# Role of Triose Phosphate Utilization in photosynthetic response of rice to variable carbon dioxide levels and plant source-sink relations

**DOI:** 10.1101/633016

**Authors:** Denis Fabre, Xinyou Yin, Michael Dingkuhn, Anne Clément-Vidal, Sandrine Roques, Lauriane Rouan, Armelle Soutiras, Delphine Luquet

## Abstract

This study aimed to understand the physiological bases of rice photosynthesis response to C source-sink imbalances, with focus on dynamics of the photosynthetic parameter TPU (Triose Phosphate Utilization). A dedicated experiment was replicated twice on IR64 indica rice cultivar in controlled environments. Plants were grown under the current ambient CO_2_ concentration until heading, thereafter, two CO_2_ treatments (400 and 800 μmol mol^−1^) were compared in the presence and absence of a panicle pruning treatment modifying the C sink. At two weeks after heading, photosynthetic parameters derived from CO_2_ response curves, and nonstructural carbohydrate content of flag leaf and internodes were measured 3-4 times of day. Spikelet number per panicle and flag leaf area on the main culm were recorded. Net C assimilation and TPU decreased progressively after midday in panicle-pruned plants, especially under 800 μmol mol^−1^. This TPU reduction was explained by sucrose accumulation in the flag leaf resulting from the sink limitation. It is suggested that TPU is involved in rice photosynthesis regulation under elevated CO_2_ conditions, and that sink limitation effects should be considered in crop models.

**Highlight:** This study provide new insights in the effect of C source-sink relationships on rice photosynthesis. TPU should be considered in photosynthesis studies under severe source-sink imbalance at elevated CO_2_.

## Introduction

Increasing world population and negative effects of global climate change on agricultural production require increased and more climate-resilient crop yields (Ainsworth, 2008; Ort et al., 2015; von Caemmerer et al., 2012). Rice (*Oryza sativa* L.) is the staple food for almost half of the population on Earth (GRiSP, 2013). To meet rice demand in 2050, its production has to increase by 2.4% annually until 2050 (Mohanty et al., 2013; Ray et al., 2013). This must be achieved in the context of climate change that is expected to have mostly negative effects on crop yields (Porter et al., 2014). But air CO_2_ elevation (e-CO_2_), expected to reach 600 to 700 μmol mol^−1^ in 2050 (IPCC 2016), will affect C_3_ crops like rice positively if efficiently used by photosynthesis. To achieve the production goal, leaf photosynthesis is a key leverage for improving crops (Evans, 2013; Lawson et al., 2012; Long et al., 2015; Ort et al., 2015), including rice (Makino, 2011; Yoshida et al., 2008; Yoshida and Horie, 2009).

A key requirement for achieving high crop productivity is to optimize carbon source-sink balance in the plants. E-CO_2_ can perturb plant carbon (C) source-sink balance as it can increase C source more than the sink (White et al., 2016), leading to leaf carbohydrate accumulation that may down-regulate photosynthesis (Burnett et al., 2016; Paul and Foyer, 2001; Shimono et al., 2010; White et al., 2016). Source-sink interactions have been intensively studied during the last two decades (Chang et al., 2017). Whether and when plant growth and production is limited by C source (chiefly, photosynthesis) or sink (demand for organ growth) is still a key research question for agronomists, plant physiologists, biochemists and crop modelers (Burnett et al., 2016).

For agronomists, this question is particularly relevant during the grain filling period (Tang et al., 2017; Wei et al., 2018; Yang and Zhang, 2010; Zhang et al., 2017). Some studies used pruning treatments to manipulate the C source (leaf) and/or sink (grains) (Cock and Yoshida, 1973; Hasegawa et al., 2013, 2016; Jing et al., 2016; Nakano et al., 1995, 2017; Shimono et al., 2010; Shinano et al., 2006), and some of these were conducted with an e-CO_2_ treatment. Conflicting results were reported regarding the response of photosynthesis to e-CO_2_ but all studies agreed that plants with larger sink capacity benefitted more from e-CO_2_.

Several physiological studies dealt with the role of non-structural carbohydrate (NSC) in C source–sink relationships under abiotic constraints such as drought (e.g. Dingkuhn et al., 2007). Experimental manipulations of plant C source and/or sink strength demonstrated that photosynthetic rate depends on C sink strength (Ainsworth and Bush, 2011; Lemoine et al., 2013; Osorio et al., 2014). Accumulation of NSC commonly occurs in leaves of plants grown under e-CO_2_ but down-regulation of photosynthesis is not always observed (Leakey et al., 2009; Wang et al., 2015), suggesting that feedbacks on photosynthesis are complex. Part of this complexity might be explained by partitioning of leaf NSC between sucrose and starch, controlled by day length (Mengin et al., 2017; Pokhilko and Ebenhoh, 2015; Sharkey, 2015; Sulpice et al., 2014), other environmental variables such as water deficit (Luquet et al., 2008), or time of day (Bläsing et al., 2005; Gibon et al., 2006). Plants lacking in sink capacity show reduced phloem loading. Rice is particularly effective in its capacity to export NSC from source leaves, suggesting that its photosynthetic response to e-CO_2_ should be efficient (Makino and Mae, 1999), but little is known on the mechanisms.

Biochemical studies on C source-sink relationships have focused on two key parameters: the utilisation of triose phosphate produced in the Calvin cycle for sucrose and starch synthesis, and ribulose biphosphate (RuBP) regeneration by inorganic phosphate (Pi) recycling, which is related to sugar turnover (Leegood and Furbank, 1986; Paul and Foyer, 2001; Paul and Pellny, 2003; Sharkey, 1985). Analysis of leaf photosynthesis classically considers three limiting steps according to a biochemical photosynthesis model (the FvCB model hereafter) described by Farquhar et al. (1980), later extended by Sharkey (1985), involving the key parameters: i) Rubisco activity (*V*_cmax_), ii) photosynthetic electron transport rate (*J*_max_) determining the ability to regenerate RuBP substrate for Rubisco, and iii) Triose Phosphate Utilization (TPU) driving the synthesis of sucrose from sugar precursors in the Calvin-Benson cycle. Thereby, TPU acts as a short-term sink that commits carbon to end-products and is closely linked to triose phosphate conversion into sucrose or starch. High sink capacity accelerates the utilization of triose phosphate for sugar synthesis and export *via* phloem. It accelerates Pi recycling and RuBP regeneration in the Calvin cycle (Gibson et al., 2011; Kant et al., 2012; Kaschuk et al., 2009; Paul and Foyer, 2001; Paul and Pellny, 2003). TPU limitation occurs primarily at high CO_2_ or sink-limited situations (Leegood and Furbank, 1986; Sharkey, 1985).

The FvCB model is commonly used as a module in crop models (Wu et al., 2016). Currently, photosynthesis is thought to be limited mainly by either *V*_cmax_ or *J*_max_, whereas TPU has received less attention and is mostly ignored by crop models (Long and Bernacchi, 2003; von Caemmerer, 2000) because its regulation is largely unknown (Yang et al., 2016). However, TPU as a link between sugar production (source) and consumption (sink) may become functionally important for crop models when addressing future climatic scenarios and e-CO_2_ (Busch and Sage, 2016), particularly in sink-limiting situations (Asseng et al., 2017; Lombardozzi et al., 2017).

As the relations between source and sink activities at the crop, plant and process levels are complex, and there is a need to integrate the different levels. For this purpose, the present study aims to explore the role of TPU in the regulation of photosynthesis in response to C source-sink relationships. A dedicated experiment was designed to observe photosynthetic parameters, the dynamics of C source-sink ratio at plant or leaf level, and NSC partitioning between soluble sugars and starch. Results are expected to provide insights on whether TPU influences photosynthesic rate in current and future climatic scenarios, and should thus be considered in crop modelling.

## Material and methods

### Plant material and growth conditions

Seeds of high yielding *indica* rice cultivar from the Philippines, IR64, were germinated on wet filter paper and transplanted to 4L pots filled with EGO 140 substrate (17%N-10%P-14%K, pH = 5). Basal fertilizer was applied using Basacot 6 M (Compo Expert) at 2 g l^−1^, 11%N-9%P-19%K +2%Mg. A second application was performed (topdressing) just before the heading stage to avoid post-floral nitrogen deficiency. Experiment was undertaken twice in the same growth chambers, in November 2016 (Exp1) and February 2017 (Exp2), using the same environmental conditions.

For each experiment, 60 plants were grown and divided between two identical growth chambers (microclima MC1750E, Snijders, Netherlands) at CIRAD, Montpellier, France. The two chambers were maintained at 12-h photoperiod, with day/night temperatures of 29/22°C, air humidity of 65/80% and daytime radiation of 1200 μmol photons m^−2^s^−1^ photosynthetically active radiation (PAR) at plant tops. The 30 pots per chamber were rotated regularly to comensate for heterogeneity. They were arranged at 35-cm plant spacing in a completely randomized design with five replicates (potted plants). Pots were irrigated to maintain soil moisture at field capacity level.

At heading stage (80 days after transplanting), all panicles of half of the plants in each growth chamber were excised (pruning treatment PR, first experimental factor). Non-PR plants were called controls. The second factor was CO_2_ treatment: In chamber 1, CO_2_ level was set at 400 μmol mol^−1^ during the whole experiment (ambient treatment); in chamber 2, CO_2_ level was maintained at 400 μmol mol^−1^ until the onset of heading, then switched to 800 μmol mol^−1^ (e-CO_2_ treatment) for 15 days, the main period of grain filling (Cho et al., 1988). At the end of the e-CO_2_ period, physiological and biochemical measurements were performed. The combination of PR and CO_2_ treatments at grain filling stage was chosen to achieve maximimal C source-sink differences and to avoid the appearance of new sinks (panicles) during differential treatments.

For Exp1 in each growth chamber, photosynthesis, biochemical and biomass measurements (see details below) were carried out at three times of day: morning, midday, and afternoon, at +1h, +6h, and +9h after dawn, respectively, on 5 consecutive days. Measurements were done on a total of 60 plants (2 PR treatments x 3 times of day of sampling x 2 e-CO_2_ levels x 5 biological replications).

For Exp2, in each growth chamber and for each treatment, photosynthesis, biochemical and biomass measurements carried out at midday, afternoon and evening; at +6h, +9h and +11h after dawn, respectively. As for Exp1, these were done for 5 consecutive days, resulting in a total of 60 measured plants.

### Leaf photosynthesis measurement

Leaf photosynthesis parameters were measured on the flag leaf on the main culm 2 weeks after heading, using two portable photosynthesis systems (GFS-3100, Walz, Germany) identically calibrated and used to measure simultaneously plants at each CO_2_ level. The measurements were made *in situ* using saturating PPFD light (1500 μmol m^−2^ s^−1^ of PAR), controlled leaf temperature at 29°C, relative humidity in the cuvette set at 65%, and constant air flow rate through the cuvette of 800 ml min^−1^. We used a large exchange area cuvette of 8 cm^2^ to limit border effects known to affect photosynthesis measurement at high [CO_2_] (Long and Bernacchi, 2003). Net photosynthesis CO_2_ response curves (*A*/*C*_i_) were obtained over a range of external CO_2_ levels in the following order: 400, 300, 200, 100, 50, 400, 600, 800, 1000, 1200, 1400, 1600, 1800 and 2000 μmol mol^−1^. At each step, gas exchange variables were recorded upon reaching steady-state (7-8 min per step, coefficient of variation <1%). In subsequent analysis, net photosynthesis (*A*), stomatal conductance (*g*_s_) and intercellular CO_2_ concentration (*C*_i_) were determined as the value measured at the 400 μmol mol^−1^ CO_2_ step of the curve. Chlorophyll fluorescence was measured for each CO_2_ step simultaneously using Walz PAM-fluorimeter 3055FL, integrated into the photosynthesis equipment. The steady-state fluorescence yield (*F*_s_) was measured after registering the gas-exchange parameters. A saturating light pulse (8000 μmol m^−2^ s^−1^ during 0.8 s) was applied to achieve the light-adapted maximum fluorescence (*F*_m_’). The operating PSII photochemical efficiency (φPSII) was determined as φPSII = (*F*_m_’ – *F*_s_)/*F*_m_’.

To fit the FvCB model of C_3_ photosynthesis to experimental data, we used non-linear fitting procedure developed by Sharkey (2016), version 2, using the Rubisco kinetic parameters determined by temperature response functions according to Bernacchi (2002). The three main photosynthesis limitations, maximum carboxylation rate (*V*_cmax_), electron transport rate (*J*_max_), and triose phosphate utilisation (TPU), were estimated simultaneously, along with mesophyll conductance (*g*_m_), by minimizing the sum of squares of the residuals. Independent measurements of day-time respiration (*R*_d_) were made on some plants using the procedure of Yin et al. (2011), and an average value of *R*_d_ was used as a constant in the fitting procedure to avoid over-parameterization. Fluorescence measurements of φPSII were used to study the rate-limiting process for each level on the CO_2_ curve, particularly to study the transition to TPU limitation as φPSII declines at high *C*_i_ (Sharkey 2016). To allow treatment comparisons, all parameters were scaled to a constant temperature of 25°C. In total, 120 CO_2_ response curves were analyzed.

### Sugar content analysis

Immediately after *A*/*C*_i_ curve measurements, the same leaf was sampled to measure non-structural carbohydrate content (NSC: starch, sucrose, glucose, fructose). Segments of the corresponding culm (top internode below the peduncle and bottom-most elongated internode) were also analyzed. Prior to grinding by ball grinder (Mixer mill MM 200, Retsch, Germany), the samples were frozen in liquid nitrogen. Sugars were extracted 3x from 20 mg samples with 1 mL of 80% ethanol for 30 min at 75°C, then centrifuged 10 min at 9500 g (Mikro 200, Hettich centrifuge). Soluble sugars (sucrose, glucose, and fructose) were contained in the supernatant and starch in the sediment. Supernatant was filtered in the presence of polyvinyl polypyrrolidone and activated carbon to eliminate pigments and polyphenols. After evaporation of solute with Speedvac (RC 1022 and RCT 90, Jouan SA, Saint Herblain, France), soluble sugars were quantified by high performance ionic chromatography (HPIC, standard Dionex) with pulsated amperometric detection (HPAE-PAD). The sediment was solubilized with 0.02 N NaOH at 90°C for 1.5 h then hydrolyzed with a-amyloglucosidase at 50°C, pH 4.2 for 1.5 h. Starch was quantified as described in Boehringer (Pomeranz and Meloan, 1994) with 5 μL of hexokinase (glucose-6-phosphate dehydrogenase), followed by photometry of NADPH at 340 nm (spectrophotometer UV/VIS V-530, Jasco Corporation, Tokyo, Japan).

### Leaf Nitrogen Content and Mass per Area

On each plant, segments of the leaf used for measuring CO_2_ curve was used for determining the nitrogen content in % dw (Nm; mg N g^−1^ dw of leaf blade) and specific leaf area (SLA; cm^2^ g^−1^). Nitrogen content per leaf area (Na; g N m^−2^) was obtained as Nm divided by SLA. The area of each sample was measured with a leaf area meter (Li-3100 Li-Cor) then oven-dried until constant weight (48 h at 70°C). Total nitrogen (N) was analyzed based by Dumas combustion method using a LECO TruMac Nitrogen analyzer, and potassium content (K) was measured in addition in Exp2 using an ICP-OES spectrometer 700 Series (Agilent Technologies). A relative indicator of chlorophyll content, SPAD, was also measured on the same leaf using a SPAD-502 (Minolta, Ltd., Japan).

### Plant growth and biomass measurements

After sampling for biochemical analyses, all the aerial parts of plants were collected. Leaf blade, sheath, culm and panicle dw per plant (DM) were measured after drying samples at 70°C during 48 h (adding *a posteriori* the DM of organ segments sampled previously). Tillers and panicles were counted and total plant green leaf area measured, using a leaf area meter (Li-3100 Li-Cor, Lincoln, NE, USA). A proxy for the source-sink ratio was estimated at the time of photosynthesis measurements as the main-culm flag leaf area to fertile spikelet number ratio.

### Statistical analysis

A three-way analysis of variance (ANOVA) of pruning treatment (PR), CO_2_, sampling time and interaction effects on each measured parameter was performed for each experiment combined using the PROC MIXED method of the SAS package (SAS Institute Inc., NC, USA, version 9.04). A multiple comparison of means and Tukey’s test (α=0,05) was then performed. Carbohydrate variables were log-transformed to stabilize variance. An analysis of covariance was performed to study the relationship between TPU, sugar contents, CO_2_ and pruning treatments, using the PROC GLIMMIX method of the SAS package. Blocking effects (time of the day) were considered as random effects (Piepho et al., 2003). No experiment effect was observed on parameters measured at the same time in both experiments (illustrated by box plots in Fig. S1 for A, TPU and flag leaf sucrose content only).

## Results

### Photosynthetic parameter responses to C source-sink imbalance

Under ambient [CO_2_], leaf photosynthesis (*A*) was significantly reduced by PR treatment (P<0.001, Table S1), mainly in the afternoon during which *A* declined for all treatments. This was supported by a significant interaction observed between pruning treatment and time of day of measurement (P<0.001, Table S1). This result was amplified under elevated [CO_2_], with a reduction of *A* by 50% in the evening for PR compared to control plants (Fig. 1A).

**Fig. 1:**
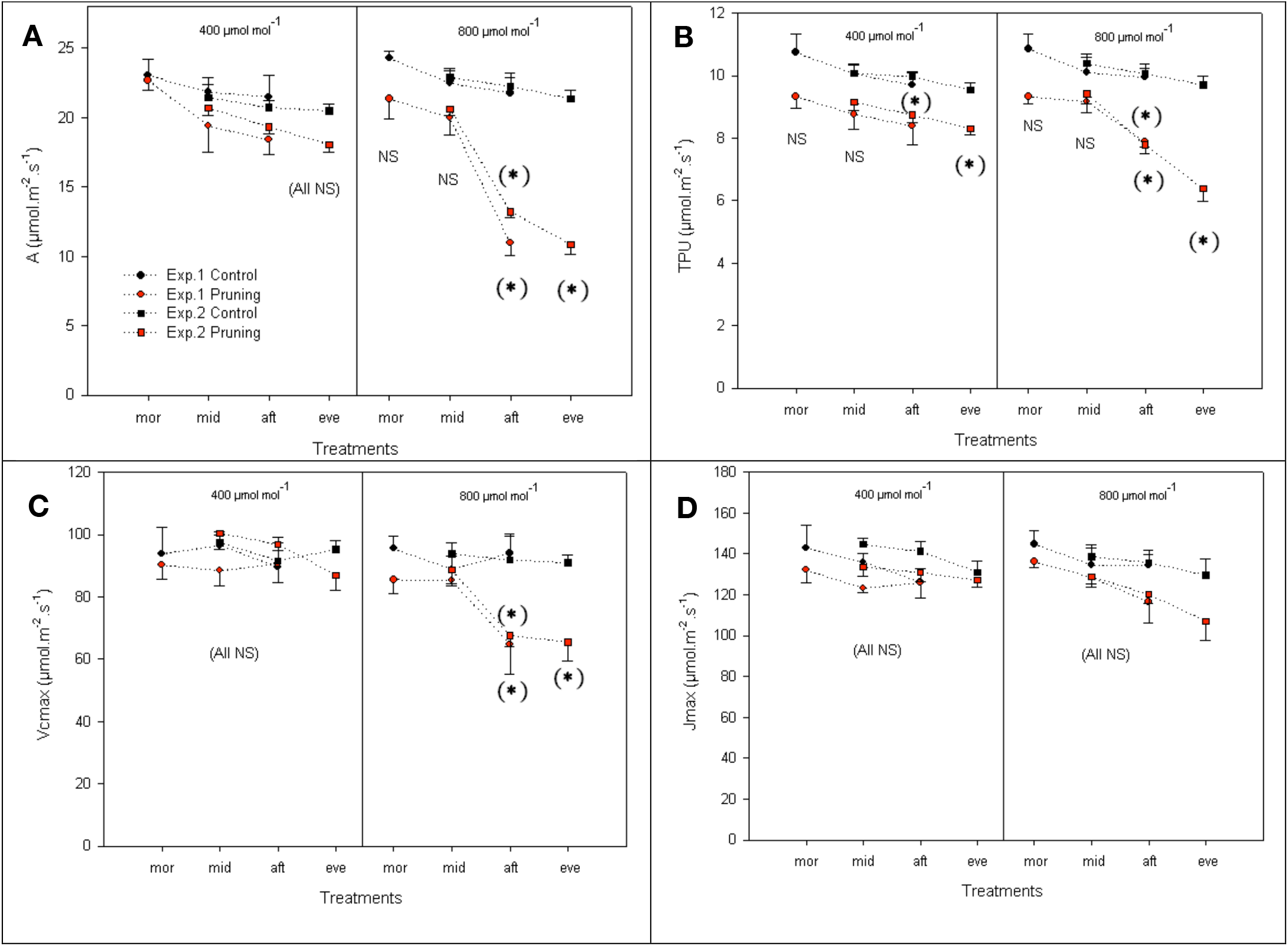
Effect of pruning and CO_2_ treatment on photosynthetic parameters, (A) net assimilation rate *A*, (B) triose phosphate utilization TPU, (C) maximum carboxylation rate of Rubisco *V*_cmax_, and (D) maximum rate of electron transport *J*_max_. Measured at 2 weeks after heading on the flag leaf on the main culm of plants of IR64 rice genotype at two CO2 levels (400 and 800 μmol mol^−1^), in two growth chambers for experiments 1 and 2. Black symbol: Control (plants with panicles) and red symbol: PR (plants with panicle pruned). Measurements were carried out at morning (mor), midday (mid), afternoon (aft) and evening (eve) periods. Stars indicate significant difference at *p* < 0.05 (Tukey HSD test) among values. NS: not significative. Each point represents the mean of 5 values ± SE.

No significant effects of the experimental factors were observed on leaf chlorophyll content (SPAD). Stomatal conductance (*g*_s_) decreased along the day at both CO_2_ concentrations, with significant effects of time of day (P<0.001, Table S1). *g*_s_ was significantly decreased by PR treatment (P<0.001) which interacted with [CO_2_] (P<0.05) without any significant variation of intercellular CO_2_ concentration (*C*_i_) (Table 1). Decrease of *g*_s_ was significant only in the afternoon under ambient [CO_2_] but already from midday onwards under elevated [CO_2_].

**Table 1:**
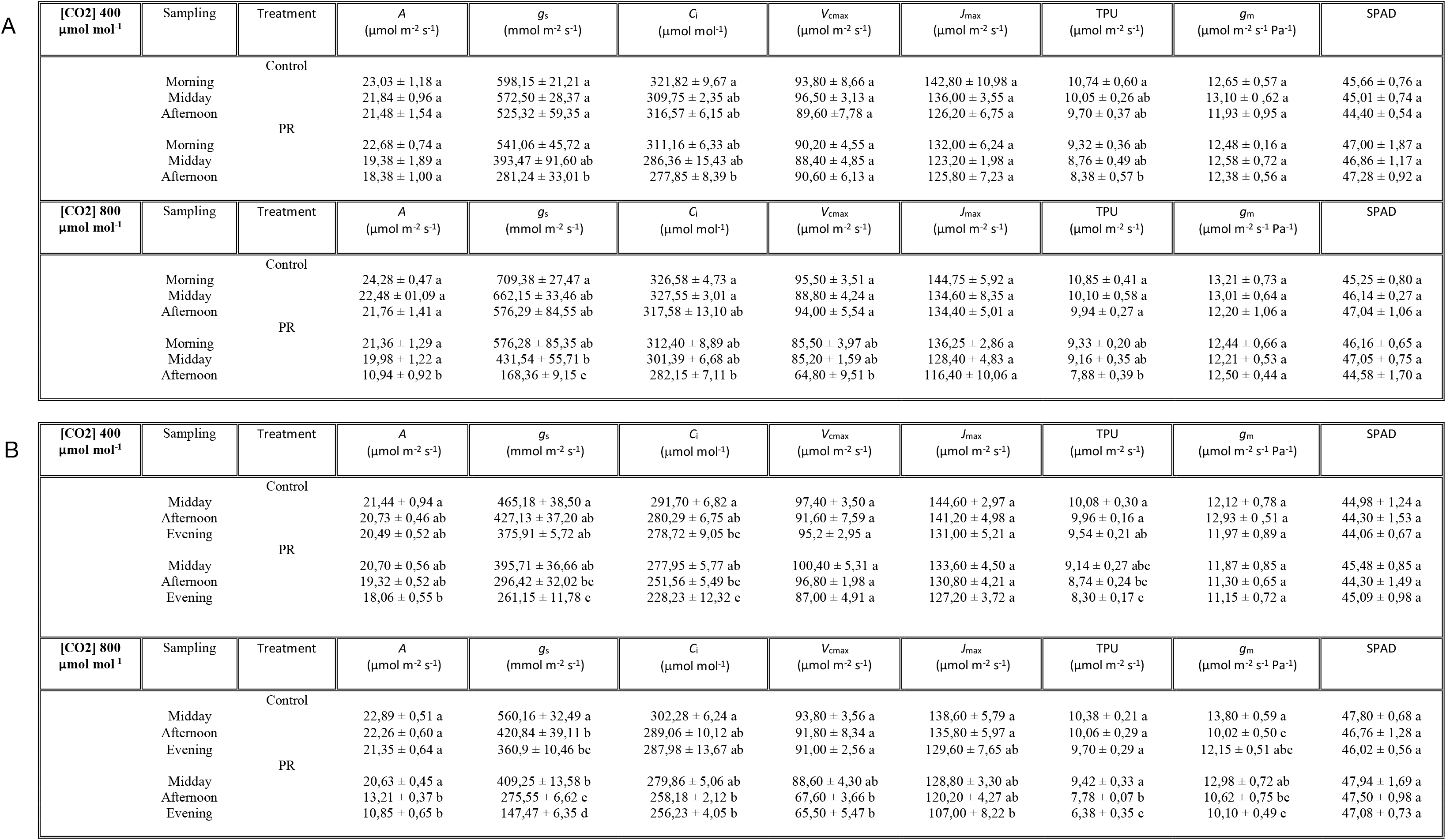
Photosynthesis characteristics (A: for Exp1 and B: for Exp2) measured two weeks after heading on the flag leaf on the main culm of IR64 plants grown under two [CO_2_] levels, with panicle pruned at heading (PR) or not (Control). Average values ± standard errors (n=5) are presented. For each column within a [CO_2_] level, values followed by different letters differ significantly (P<0.05)

| [CO<sub>2</sub>] 400 $\mu\text{mol mol}^{-1}$ | Sampling | Treatment | A<br>($\mu\text{mol m}^{-2} \text{s}^{-1}$) | $g_s$<br>($\text{mmol m}^{-2} \text{s}^{-1}$) | $C_i$<br>($\mu\text{mol mol}^{-1}$) | $V_{\text{cmax}}$<br>($\mu\text{mol m}^{-2} \text{s}^{-1}$) | $J_{\text{max}}$<br>($\mu\text{mol m}^{-2} \text{s}^{-1}$) | TPU<br>($\mu\text{mol m}^{-2} \text{s}^{-1}$) | $g_m$<br>($\mu\text{mol m}^{-2} \text{s}^{-1} \text{Pa}^{-1}$) | SPAD |
| --- | --- | --- | --- | --- | --- | --- | --- | --- | --- | --- |
| Control |  |  |  |  |  |  |  |  |  |  |
| | Morning | | 23,03 ± 1,18 a | 598,15 ± 21,21 a | 321,82 ± 9,67 a | 93,80 ± 8,66 a | 142,80 ± 10,98 a | 10,74 ± 0,60 a | 12,65 ± 0,57 a | 45,66 ± 0,76 a |
| | Midday | | 21,84 ± 0,96 a | 572,50 ± 28,37 a | 309,75 ± 2,35 ab | 96,50 ± 3,13 a | 136,00 ± 3,55 a | 10,05 ± 0,26 ab | 13,10 ± 0,62 a | 45,01 ± 0,74 a |
| | Afternoon | | 21,48 ± 1,54 a | 525,32 ± 59,35 a | 316,57 ± 6,15 ab | 89,60 ± 7,78 a | 126,20 ± 6,75 a | 9,70 ± 0,37 ab | 11,93 ± 0,95 a | 44,40 ± 0,54 a |
| PR |  |  |  |  |  |  |  |  |  |  |
| | Morning | | 22,68 ± 0,74 a | 541,06 ± 45,72 a | 311,16 ± 6,33 ab | 90,20 ± 4,55 a | 132,00 ± 6,24 a | 9,32 ± 0,36 ab | 12,48 ± 0,16 a | 47,00 ± 1,87 a |
| | Midday | | 19,38 ± 1,89 a | 393,47 ± 91,60 ab | 286,36 ± 15,43 ab | 88,40 ± 4,85 a | 123,20 ± 1,98 a | 8,76 ± 0,49 ab | 12,58 ± 0,72 a | 46,86 ± 1,17 a |
| | Afternoon | | 18,38 ± 1,00 a | 281,24 ± 33,01 b | 277,85 ± 8,39 b | 90,60 ± 6,13 a | 125,80 ± 7,23 a | 8,38 ± 0,57 b | 12,38 ± 0,56 a | 47,28 ± 0,92 a |
| [CO<sub>2</sub>] 800 $\mu\text{mol mol}^{-1}$ | Sampling | Treatment | A<br>($\mu\text{mol m}^{-2} \text{s}^{-1}$) | $g_s$<br>($\text{mmol m}^{-2} \text{s}^{-1}$) | $C_i$<br>($\mu\text{mol mol}^{-1}$) | $V_{\text{cmax}}$<br>($\mu\text{mol m}^{-2} \text{s}^{-1}$) | $J_{\text{max}}$<br>($\mu\text{mol m}^{-2} \text{s}^{-1}$) | TPU<br>($\mu\text{mol m}^{-2} \text{s}^{-1}$) | $g_m$<br>($\mu\text{mol m}^{-2} \text{s}^{-1} \text{Pa}^{-1}$) | SPAD |
| Control |  |  |  |  |  |  |  |  |  |  |
| | Morning | | 24,28 ± 0,47 a | 709,38 ± 27,47 a | 326,58 ± 4,73 a | 95,50 ± 3,51 a | 144,75 ± 5,92 a | 10,85 ± 0,41 a | 13,21 ± 0,73 a | 45,25 ± 0,80 a |
| | Midday | | 22,48 ± 01,09 a | 662,15 ± 33,46 ab | 327,55 ± 3,01 a | 88,80 ± 4,24 a | 134,60 ± 8,35 a | 10,10 ± 0,58 a | 13,01 ± 0,64 a | 46,14 ± 0,27 a |
| | Afternoon | | 21,76 ± 1,41 a | 576,29 ± 84,55 ab | 317,58 ± 13,10 ab | 94,00 ± 5,54 a | 134,40 ± 5,01 a | 9,94 ± 0,27 a | 12,20 ± 1,06 a | 47,04 ± 1,06 a |
| PR |  |  |  |  |  |  |  |  |  |  |
| | Morning | | 21,36 ± 1,29 a | 576,28 ± 85,35 ab | 312,40 ± 8,89 ab | 85,50 ± 3,97 ab | 136,25 ± 2,86 a | 9,33 ± 0,20 ab | 12,44 ± 0,66 a | 46,16 ± 0,65 a |
| | Midday | | 19,98 ± 1,22 a | 431,54 ± 55,71 b | 301,39 ± 6,68 ab | 85,20 ± 1,59 ab | 128,40 ± 4,83 a | 9,16 ± 0,35 ab | 12,21 ± 0,53 a | 47,05 ± 0,75 a |
| | Afternoon | | 10,94 ± 0,92 b | 168,36 ± 9,15 c | 282,15 ± 7,11 b | 64,80 ± 9,51 b | 116,40 ± 10,06 a | 7,88 ± 0,39 b | 12,50 ± 0,44 a | 44,58 ± 1,70 a |

B
| [CO<sub>2</sub>] 400 $\mu\text{mol mol}^{-1}$ | Sampling | Treatment | A<br>($\mu\text{mol m}^{-2} \text{s}^{-1}$) | $g_s$<br>($\text{mmol m}^{-2} \text{s}^{-1}$) | $C_i$<br>($\mu\text{mol mol}^{-1}$) | $V_{\text{cmax}}$<br>($\mu\text{mol m}^{-2} \text{s}^{-1}$) | $J_{\text{max}}$<br>($\mu\text{mol m}^{-2} \text{s}^{-1}$) | TPU<br>($\mu\text{mol m}^{-2} \text{s}^{-1}$) | $g_m$<br>($\mu\text{mol m}^{-2} \text{s}^{-1} \text{Pa}^{-1}$) | SPAD |
| --- | --- | --- | --- | --- | --- | --- | --- | --- | --- | --- |
| Control |  |  |  |  |  |  |  |  |  |  |
| | Midday | | 21,44 ± 0,94 a | 465,18 ± 38,50 a | 291,70 ± 6,82 a | 97,40 ± 3,50 a | 144,60 ± 2,97 a | 10,08 ± 0,30 a | 12,12 ± 0,78 a | 44,98 ± 1,24 a |
| | Afternoon | | 20,73 ± 0,46 ab | 427,13 ± 37,20 ab | 280,29 ± 6,75 ab | 91,60 ± 7,59 a | 141,20 ± 4,98 a | 9,96 ± 0,16 a | 12,93 ± 0,51 a | 44,30 ± 1,53 a |
| | Evening | | 20,49 ± 0,52 ab | 375,91 ± 5,72 ab | 278,72 ± 9,05 bc | 95,2 ± 2,95 a | 131,00 ± 5,21 a | 9,54 ± 0,21 ab | 11,97 ± 0,89 a | 44,06 ± 0,67 a |
| PR |  |  |  |  |  |  |  |  |  |  |
| | Midday | | 20,70 ± 0,56 ab | 395,71 ± 36,66 ab | 277,95 ± 5,77 ab | 100,40 ± 5,31 a | 133,60 ± 4,50 a | 9,14 ± 0,27 abc | 11,87 ± 0,85 a | 45,48 ± 0,85 a |
| | Afternoon | | 19,32 ± 0,52 ab | 296,42 ± 32,02 bc | 251,56 ± 5,49 bc | 96,80 ± 1,98 a | 130,80 ± 4,21 a | 8,74 ± 0,24 bc | 11,30 ± 0,65 a | 44,30 ± 1,49 a |
| | Evening | | 18,06 ± 0,55 b | 261,15 ± 11,78 c | 228,23 ± 12,32 c | 87,00 ± 4,91 a | 127,20 ± 3,72 a | 8,30 ± 0,17 c | 11,15 ± 0,72 a | 45,09 ± 0,98 a |
| [CO<sub>2</sub>] 800 $\mu\text{mol mol}^{-1}$ | Sampling | Treatment | A<br>($\mu\text{mol m}^{-2} \text{s}^{-1}$) | $g_s$<br>($\text{mmol m}^{-2} \text{s}^{-1}$) | $C_i$<br>($\mu\text{mol mol}^{-1}$) | $V_{\text{cmax}}$<br>($\mu\text{mol m}^{-2} \text{s}^{-1}$) | $J_{\text{max}}$<br>($\mu\text{mol m}^{-2} \text{s}^{-1}$) | TPU<br>($\mu\text{mol m}^{-2} \text{s}^{-1}$) | $g_m$<br>($\mu\text{mol m}^{-2} \text{s}^{-1} \text{Pa}^{-1}$) | SPAD |
| Control |  |  |  |  |  |  |  |  |  |  |
| | Midday | | 22,89 ± 0,51 a | 560,16 ± 32,49 a | 302,28 ± 6,24 a | 93,80 ± 3,56 a | 138,60 ± 5,79 a | 10,38 ± 0,21 a | 13,80 ± 0,59 a | 47,80 ± 0,68 a |
| | Afternoon | | 22,26 ± 0,60 a | 420,84 ± 39,11 b | 289,06 ± 10,12 ab | 91,80 ± 8,34 a | 135,80 ± 5,97 a | 10,06 ± 0,29 a | 10,02 ± 0,50 c | 46,76 ± 1,28 a |
| | Evening | | 21,35 ± 0,64 a | 360,9 ± 10,46 bc | 287,98 ± 13,67 ab | 91,00 ± 2,56 a | 129,60 ± 7,65 ab | 9,70 ± 0,29 a | 12,15 ± 0,51 abc | 46,02 ± 0,56 a |
| PR |  |  |  |  |  |  |  |  |  |  |
| | Midday | | 20,63 ± 0,45 a | 409,25 ± 13,58 b | 279,86 ± 5,06 ab | 88,60 ± 4,30 ab | 128,80 ± 3,30 ab | 9,42 ± 0,33 a | 12,98 ± 0,72 ab | 47,94 ± 1,69 a |
| | Afternoon | | 13,21 ± 0,37 b | 275,55 ± 6,62 c | 258,18 ± 2,12 b | 67,60 ± 3,66 b | 120,20 ± 4,27 ab | 7,78 ± 0,07 b | 10,62 ± 0,75 bc | 47,50 ± 0,98 a |
| | Evening | | 10,85 ± 0,65 b | 147,47 ± 6,35 d | 256,23 ± 4,05 b | 65,50 ± 5,47 b | 107,00 ± 8,22 b | 6,38 ± 0,35 c | 10,10 ± 0,49 c | 47,08 ± 0,73 a |

Before estimating *g*_m_ and other derived parameters from measured *A*/*C*_i_ curves, we assessed the shape of *A*/*C*_i_ response curves (replicate means) for each treatment along the day (Fig. 2A). Within high *C*_i_ levels, *A* responded little to a change in *C*_i_. It even declined with increasing *C*_i_ for the PR-treated plants (Fig. 2A), suggesting an inhibition of *A* by TPU limitation. It is difficult to rely on only *A*/*C*_i_ curves to determine the transition from RuBP-regeneration limitation to TPU limitation since they usually occur together under high [CO_2_] (Bernacchi et al., 2013; Long and Bernacchi, 2003). We used chlorophyll fluorescence-based data on the operating efficiency of PSII electron flow (φPSII) measured concomitantly with *A* to detect the *C*_i_ above which TPU limited *A*. When this is the case, φPSII declines (Sharkey, 2016). The decline of φPSII was observed above a *C*_i_ of 825 μmol mol^−1^ in the evening, in plants exposed to ambient [CO_2_] and pruning treatment (Fig. 2B, left). It occurred at *C*_i_ above 742 and 363 μmol mol^−1^ under elevated CO_2_ condition in the afternoon and in the evening, respectively (Fig. 2B, right). As these *C*_i_ thresholds were mostly higher than the *C*_i_ at *A* measurement (Fig. 2B), TPU did not limit *A* under the experimental conditions. However, under the most severe sink limitation (PR, 800 μmol mol^−1^ [CO_2_], evening) TPU was close to limiting levels.

**Fig. 2:**
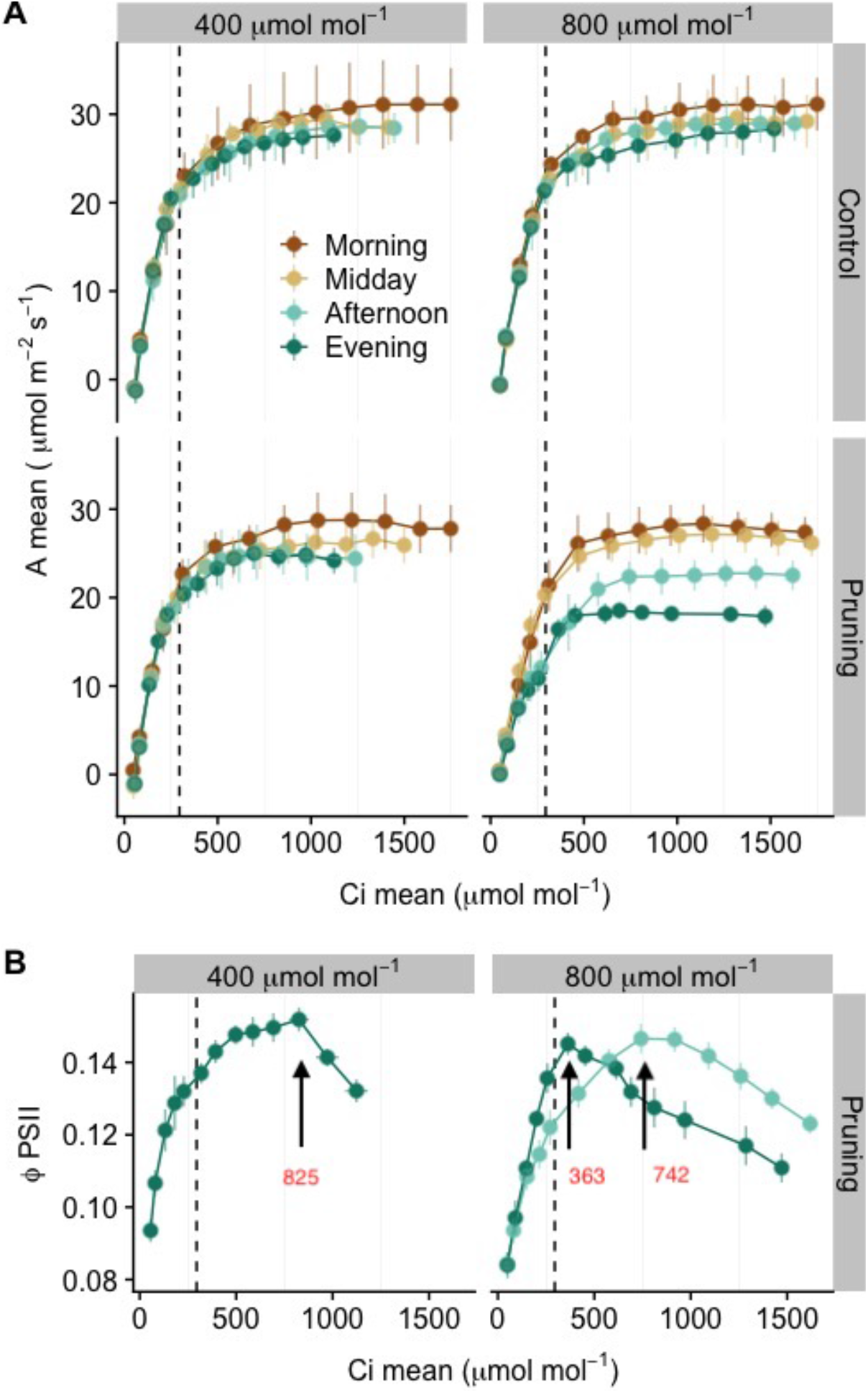
Mean *A*/*C*_i_ curves for all treatment combination (A) and corresponding mean φPSII/*C*_i_ curves (B) for experiments 1 and 2. (shown only for Pruning treatment which caused significant TPU decline). Dashed lines indicate mean *C*_i_ value (290 μmol mol^−1^) for photosynthesis measurement at treatment CO_2_ level (400 or 800 μmol mol^−1^). Arrows in Fig. 2B indicates the *C*_i_ level at which TPU limitation may begin to occur. The CO_2_ concentrations indicated in Figure headers are treatment conditions and not those administered when measuring CO_2_ response, which comprised 14 different levels.

We applied the procedure of Sharkey (2016) to fit each *A*/*C*_i_ curve to derive estimates of *g*_m_ and biochemical photosynthetic parameters. All values of *g*_m_ were high, > 10 μmol m^−2^ s^−1^ Pa^−1^ (Table 1), values known not to limit photosynthesis. Therefore, our estimates of *g*_m_ did not explain differences in *A* among treatments or times of day.

A significant decrease was observed for *V*_cmax_ in response to PR treatment (for both experiments: P<0.001, Table S1). However, this reduction depended on CO_2_ treatment (interaction PR x CO_2_ at P<0.05, Table S1) and was significant only in the afternoon and evening under elevated CO_2_ (Table 1). Mean reduction in PR was 29% compared to control in the afternoon under elevated CO_2_ (Fig. 1C). Regarding *J*_max_, a significant effect of PR treatment was observed (P<0.001, Table S1), despite no significative numerical decrease of *J*_max_ in PR treatment compared to control as shown in Table 1 and Fig. 1D. A time-of-day effect was also observed (P<0.05).

Although *A*/*C*_i_ curves for control plants grown at 400 μmol mol^−1^ [CO_2_] did not show TPU limitation (Fig. 2), TPU was significantly reduced by PR treatment (P<0.001, Table S1). A decline of TPU after noon was observed in both [CO_2_] treatments (Fig. 1B), resulting in a highly significant time-of-day effect (P<0.001). This decrease was particularly strong under e-CO_2_ when combined with PR treatment, which led to significant interaction effects (PR x CO_2_, P<0.05). In this latter situation, significant differences between control and PR plants were observed in the afternoon. The decrease of TPU caused by PR in the evening was 40% under elevated [CO_2_] and 13% under ambient [CO_2_] (Fig.1B).

### Nonstructural carbohydrate response to C source-sink imbalance

Leaf sucrose concentration in PR plants was significantly higher than in control plants in the afternoon (P<0.001 for both PR and time-of-day effects, Table S2). No interaction effects between these factors were observed. The [CO_2_] effect on leaf sucrose was smaller (P<0.05). Hexose concentration in the flag leaf was not affected by any of the experimental factors.

PR reduced sucrose concentration in the basal internode on the main culm (P<0.001, Tables 2 and S2), without significant variations along the day. Similar results were observed for hexose concentration in the lower internode, but at about 35-fold lower concentrations than sucrose (Table 2). As soluble sugar content (hexose and sucrose) was similar in basal and upper internodes, results are presented only for basal internodes.

**Table 2:** Nonstructural carbohydrate contents, raw data (A: for Exp1 and B: for Exp2) measured two weeks after heading in the flag leaf, the upper and lower internode on the main culm of IR64 plants grown under two air CO_2_ levels, with panicle pruned at heading (PR) or not (Control). Average values ± standard errors (n=5) are presented. For each column within a [CO_2_] level, values followed by different letters differ significantly (P<0.05).

| [CO<sub>2</sub>] 400 $\mu\text{mol mol}^{-1}$ | Sampling | Treatment | Sucrose flag leaf ($\mu\text{g cm}^{-2}$) | Sucrose lower internode ($\text{mg g}^{-1}$ DM) | Hexose flag leaf ($\mu\text{g cm}^{-2}$) | Hexose lower internode ($\text{mg g}^{-1}$ DM) | Starch flag leaf ($\mu\text{g cm}^{-2}$) | Starch upper internode ($\text{mg g}^{-1}$ DM) | Starch lower internode ($\text{mg g}^{-1}$ DM) |
| --- | --- | --- | --- | --- | --- | --- | --- | --- | --- |
|  | Control |  |  |  |  |  |  |  |  |
| | Morning | | 191,17 ± 23,57 b | 150,67 ± 37,61 a | 16,02 ± 4,95 a | 11,65 ± 4,40 a | 83,07 ± 35,01 a | 134,76 ± 44,89 b | 245,23 ± 60,21 b |
| | Midday | | 234,21 ± 45,89 ab | 205,92 ± 15,09 a | 28,76 ± 6,61 a | 5,27 ± 0,79 ab | 125,66 ± 19,73 a | 187,78 ± 23,86 ab | 275,73 ± 67,09 ab |
| | Afternoon | | 270,82 ± 24,87 ab | 190,22 ± 38,72 a | 25,75 ± 3,81 a | 5,36 ± 1,01 ab | 82,85 ± 12,73 a | 163,16 ± 14,42 ab | 233,97 ± 70,05 b |
|  | PR |  |  |  |  |  |  |  |  |
| | Morning | | 231,81 ± 36,16 b | 103,53 ± 23,33 a | 20,48 ± 6,40 a | 2,19 ± 0,50 b | 84,39 ± 31,10 a | 311,88 ± 30,32 a | 519,13 ± 22,72 a |
| | Midday | | 286,91 ± 28,58 ab | 116,68 ± 7,17 a | 26,50 ± 5,43 a | 2,27 ± 0,29 b | 53,02 ± 11,54 a | 311,50 ± 38,84 a | 424,59 ± 16,60 ab |
| | Afternoon | | 395,74 ± 34,50 a | 116,83 ± 35,88 a | 26,32 ± 4,18 a | 2,93 ± 0,53 b | 127,95 ± 35,12 a | 323,19 ± 46,70 a | 531,41 ± 44,68 a |
| [CO<sub>2</sub>] 800 $\mu\text{mol mol}^{-1}$ | Sampling | Treatment | Sucrose flag leaf ($\mu\text{g cm}^{-2}$) | Sucrose lower internode ($\text{mg g}^{-1}$ DM) | Hexose flag leaf ($\mu\text{g cm}^{-2}$) | Hexose lower internode ($\text{mg g}^{-1}$ DM) | Starch flag leaf ($\mu\text{g cm}^{-2}$) | Starch upper internode ($\text{mg g}^{-1}$ DM) | Starch lower internode ($\text{mg g}^{-1}$ DM) |
|  | Control |  |  |  |  |  |  |  |  |
| | Morning | | 211,07 ± 17,99 d | 162,40 ± 26,30 a | 20,14 ± 4,95 a | 5,39 ± 1,25 a | 118,68 ± 57,70 a | 466,75 ± 26,74 a | 402,95 ± 87,42 a |
| | Midday | | 275,96 ± 20,25 cd | 134,89 ± 32,73 a | 35,35 ± 9,28 a | 5,87 ± 2,80 a | 140,17 ± 27,54 a | 402,87 ± 83,40 a | 361,65 ± 83,60 a |
| | Afternoon | | 374,44 ± 17,42 ab | 109,29 ± 31,38 a | 28,02 ± 2,32 a | 3,73 ± 0,96 a | 299,80 ± 69,52 a | 403,31 ± 38,06 a | 466,65 ± 55,21 a |
|  | PR |  |  |  |  |  |  |  |  |
| | Morning | | 289,59 ± 35,19 bcd | 77,14 ± 36,31 a | 28,59 ± 2,77 a | 1,86 ± 0,77 a | 231,67 ± 78,24 a | 476,33 ± 71,84 a | 580,20 ± 59,77 a |
| | Midday | | 361,39 ± 13,51 bc | 81,31 ± 25,90 a | 37,80 ± 5,49 a | 1,82 ± 0,45 a | 309,85 ± 85,62 a | 357,17 ± 37,35 a | 530,03 ± 40,33 a |
| | Afternoon | | 466,86 ± 24,82 a | 80,86 ± 9,98 a | 38,27 ± 1,63 a | 2,17 ± 0,28 a | 249,24 ± 69,02 a | 511,71 ± 56,08 a | 641,20 ± 26,05 a |

B
| [CO<sub>2</sub>] 400 $\mu\text{mol mol}^{-1}$ | Sampling | Treatment | Sucrose flag leaf ($\mu\text{g cm}^{-2}$) | Sucrose lower internode ($\text{mg g}^{-1}$ DM) | Hexose flag leaf ($\mu\text{g cm}^{-2}$) | Hexose lower internode ($\text{mg g}^{-1}$ DM) | Starch flag leaf ($\mu\text{g cm}^{-2}$) | Starch upper internode ($\text{mg g}^{-1}$ DM) | Starch lower internode ($\text{mg g}^{-1}$ DM) |
| --- | --- | --- | --- | --- | --- | --- | --- | --- | --- |
|  | Control |  |  |  |  |  |  |  |  |
| | Midday | | 310,98 ± 55,11 b | 161,37 ± 10,79 a | 22,34 ± 5,64 a | 12,99 ± 3,85 ab | 44,20 ± 8,28 a | 170,36 ± 47,47 ab | 156,27 ± 41,56 a |
| | Afternoon | | 355,45 ± 28,85 b | 180,03 ± 14,49 a | 18,05 ± 1,10 a | 14,71 ± 2,35 a | 70,08 ± 10,64 a | 93,63 ± 15,00 b | 178,80 ± 35,69 a |
| | Evening | | 430,43 ± 59,40 ab | 132,44 ± 12,12 a | 21,55 ± 3,91 a | 13,32 ± 0,83 b | 117,66 ± 31,14 a | 157,34 ± 55,66 ab | 147,55 ± 65,72 a |
|  | PR |  |  |  |  |  |  |  |  |
| | Midday | | 311,76 ± 33,78 b | 139,35 ± 13,67 a | 17,17 ± 1,13 a | 8,97 ± 2,03 ab | 48,51 ± 8,99 a | 170,59 ± 32,28 ab | 362,25 ± 51,80 a |
| | Afternoon | | 490,53 ± 23,50 ab | 141,84 ± 9,84 a | 26,10 ± 1,42 a | 8,00 ± 2,08 ab | 94,55 ± 8,97 a | 202,02 ± 19,43 ab | 288,21 ± 68,60 a |
| | Evening | | 496,92 ± 37,84 a | 134,44 ± 20,75 a | 28,81 ± 3,65 a | 5,72 ± 1,57 ab | 128,38 ± 33,51 a | 299,98 ± 53,68 a | 294,26 ± 63,43 a |
| [CO<sub>2</sub>] 800 $\mu\text{mol mol}^{-1}$ | Sampling | Treatment | Sucrose flag leaf ($\mu\text{g cm}^{-2}$) | Sucrose lower internode ($\text{mg g}^{-1}$ DM) | Hexose flag leaf ($\mu\text{g cm}^{-2}$) | Hexose lower internode ($\text{mg g}^{-1}$ DM) | Starch flag leaf ($\mu\text{g cm}^{-2}$) | Starch upper internode ($\text{mg g}^{-1}$ DM) | Starch lower internode ($\text{mg g}^{-1}$ DM) |
|  | Control |  |  |  |  |  |  |  |  |
| | Midday | | 267,33 ± 27,60 b | 105,41 ± 3,38 a | 20,17 ± 2,72 a | 8,53 ± 4,35 a | 53,16 ± 15,27 a | 110,36 ± 16,20 b | 184,22 ± 55,74 a |
| | Afternoon | | 352,10 ± 40,06 ab | 93,10 ± 3,78 a | 26,24 ± 4,07 a | 9,40 ± 2,24 a | 104,30 ± 17,14 a | 54,03 ± 7,15 b | 168,88 ± 60,77 a |
| | Evening | | 393,25 ± 18,79 a | 106,22 ± 8,73 a | 20 ± 2,31 a | 3,72 ± 1,07 a | 149,94 ± 23,65 a | 80,99 ± 16,68 b | 164,39 ± 52,57 a |
|  | PR |  |  |  |  |  |  |  |  |
| | Midday | | 382,65 ± 16,76 ab | 97,01 ± 17,13 a | 24,47 ± 3,23 a | 4,23 ± 0,83 a | 50,82 ± 11,22 a | 282,52 ± 32,88 a | 352,03 ± 49,52 a |
| | Afternoon | | 423,82 ± 33,98 a | 85,91 ± 9,68 a | 24,15 ± 4,96 a | 2,35 ± 0,16 a | 104,30 ± 32,82 a | 240,88 ± 31,66 a | 338,46 ± 37,24 a |
| | Evening | | 452,95 ± 12,27 a | 96,18 ± 5,58 a | 23,70 ± 3,51 a | 3,84 ± 0,62 a | 131,61 ± 39,77 a | 267,47 ± 30,55 a | 323,30 ± 37,70 a |

PR increased starch concentration in both top and bottom internodes on the main culm (P<0.001; Tables 2 and S2), whereby no interaction between CO_2_ and PR treatments was observed. No time-of-day effect was observed for starch concentration in internodes.

No PR and CO_2_ effects were observed on leaf starch concentration, but there was a significant time-of-day effect (P<0.001, Tables 2 and S2), causing a continuous increase of leaf starch concentration along the day (Table 2).

### Plant growth response to C source-sink imbalance

PR significantly increased culm dry matter (by 50-60%, P<0.001, Table S3 and Table S4) and sheath dry matter (by 12-20%, P<0.001). No [CO_2_] effect and no interaction between factors were observed (Table S3). Panicle dry weight sampled two weeks after heading in the control plants was 280% higher under elevated [CO_2_] compared to ambient [CO_2_] (P<0.001, Table S3 and Table S4), suggesting a strong stimulation of CO_2_ enrichment on grain filling. By contrast, none of the factors affected plant total leaf dry matter, tiller number, panicle number and the SLA of the flag leaves used for photosynthesis measurement (Table S3). The same was true for nitrogen and potassium contents of the flag leaf, except for a significant reduction (P<0.05) of nitrogen content under elevated [CO_2_], particularly on control plants (Table S3 and S4). Dry matter of plant roots was also measured at the end of the experiments. No significant effect of experimental factors were observed (data not presented).

### Correlations between photosynthetic and biochemical parameters

A positive linear correlation was observed between *A* and TPU (R^2^=0.64, P<0.001) across all combinations of [CO_2_] and pruning treatments, and across all times of day (Fig. 3). The strongest treatment-specific correlation was observed when PR treatment was combined with high [CO_2_], i.e. for the treatment combination causing the highest C source-sink ratio (R^2^=0.72, P<0.001). The corresponding correlation between *A* and *V*_cmax_ was also significant but weaker (R^2^=0.60; data not presented).

**Fig. 3:**
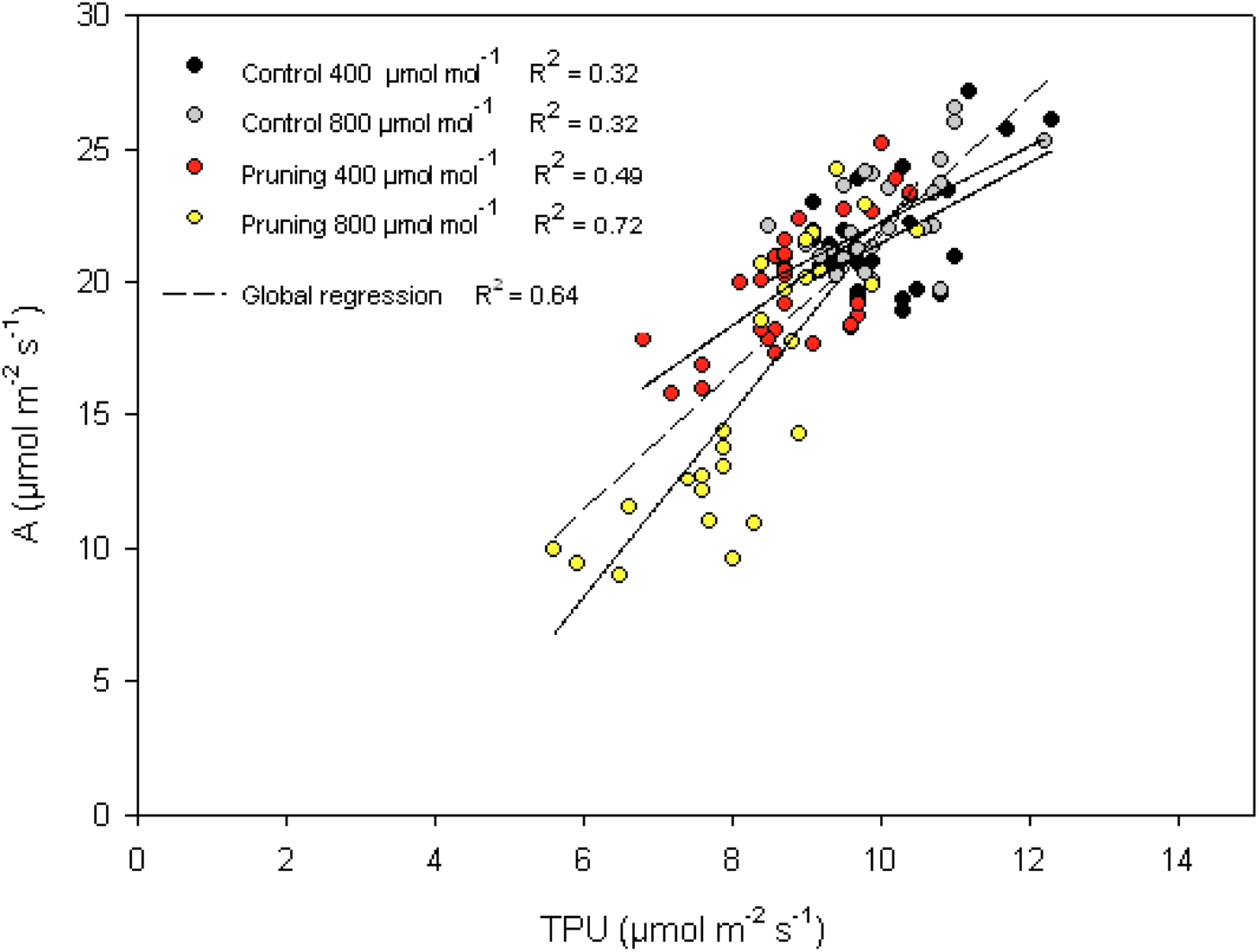
Relationship between TPU and net photosynthesis (*A*) within experiments 1 and 2 and for each combination of CO_2_ x panicle pruning treatment. With panicle and 400 μmol mol^−1^ (black symbol), with panicle and 800 μmol mol^−1^ (grey symbol), Panicles pruned and 400 μmol mol^−1^ (red symbol), panicles pruned and 800 μmol mol^−1^ (yellow symbol). Each point represents a single value.

Analysis of covariance was performed to study the relationship between nonstructural carbohydrate and TPU variations. Flag leaf sucrose concentration was by far the most predictive factor of TPU variation (P<0.001) (Table S5). This was supported by the negative, linear correlations (R^2^=0.66 for controls, R^2^=0.40 for PR) observed between flag leaf sucrose concentration and TPU (Fig. 4). The two linear correlations showed a similar slope (−5.4 for control and −6.1 for pruned) but with lower TPU value at the intercepts in the case of pruned plants. An effect of starch concentration in the lower internodes was also observed (P=0.01) but it was smaller than that of leaf sucrose.

**Fig. 4:**
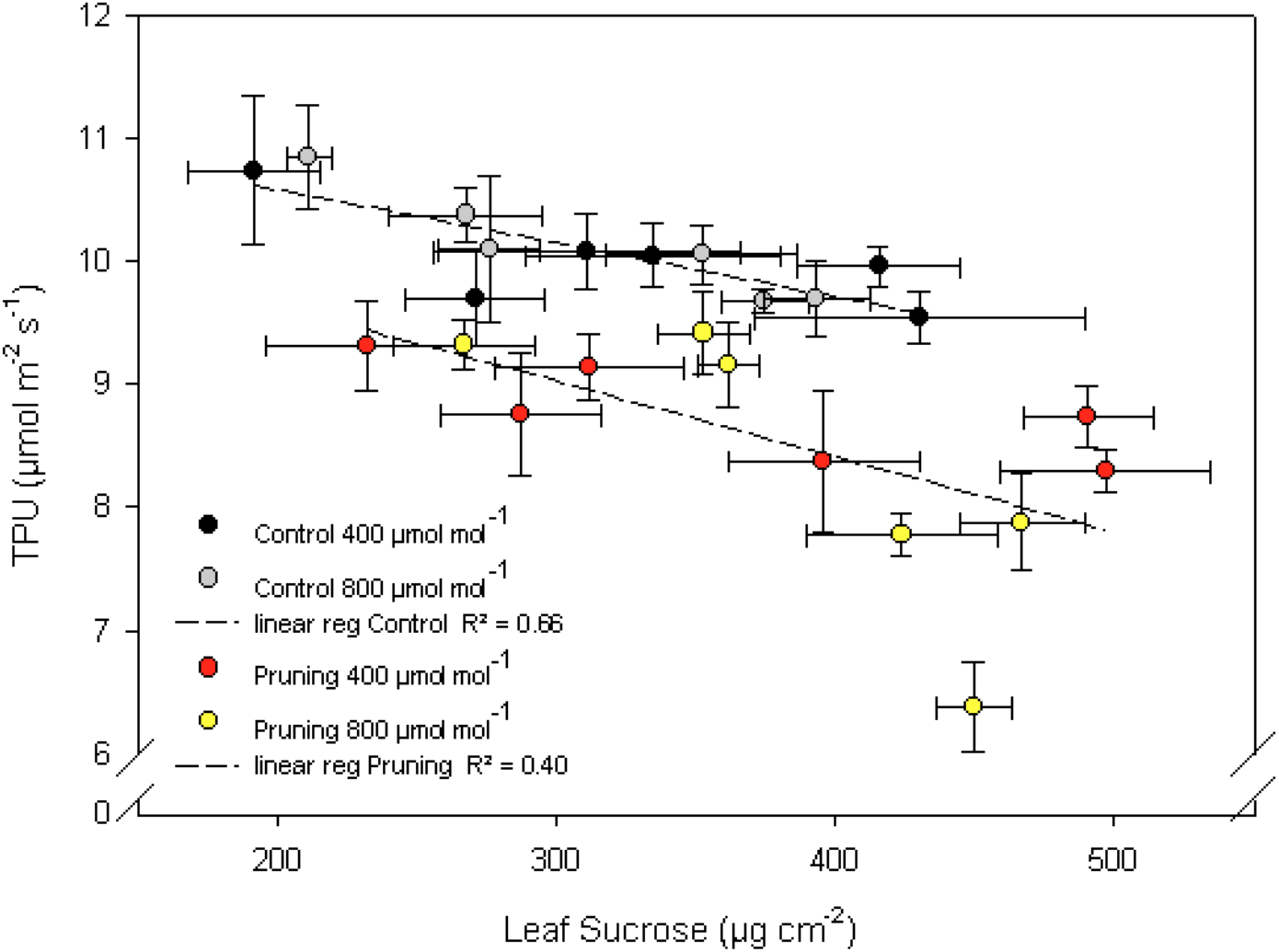
Relationship between TPU and Leaf sucrose content in experiments 1 and 2, separating control and panicle pruning treatment. With panicle at 400 μmol mol^−1^ (black symbol), with panicle at 800 μmol mol^−1^ (grey symbol), Panicles pruned at 400 μmol mol^−1^ (red symbol), panicles pruned at 800 μmol mol^−1^ (yellow symbol). Dashed lines represent linear regression for both pruning treatments. Each point is the average of 5 values and is presented with horizontal and vertical standard errors.

Finally, a negative correlation was found between TPU and plant C source-sink ratio measured two weeks after heading (R^2^ of 0.45, P<0.01; Fig. 5), defined as the ratio of flag leaf area over fertile spikelet number of the corresponding panicle (measured only for Exp2).

**Fig. 5:**
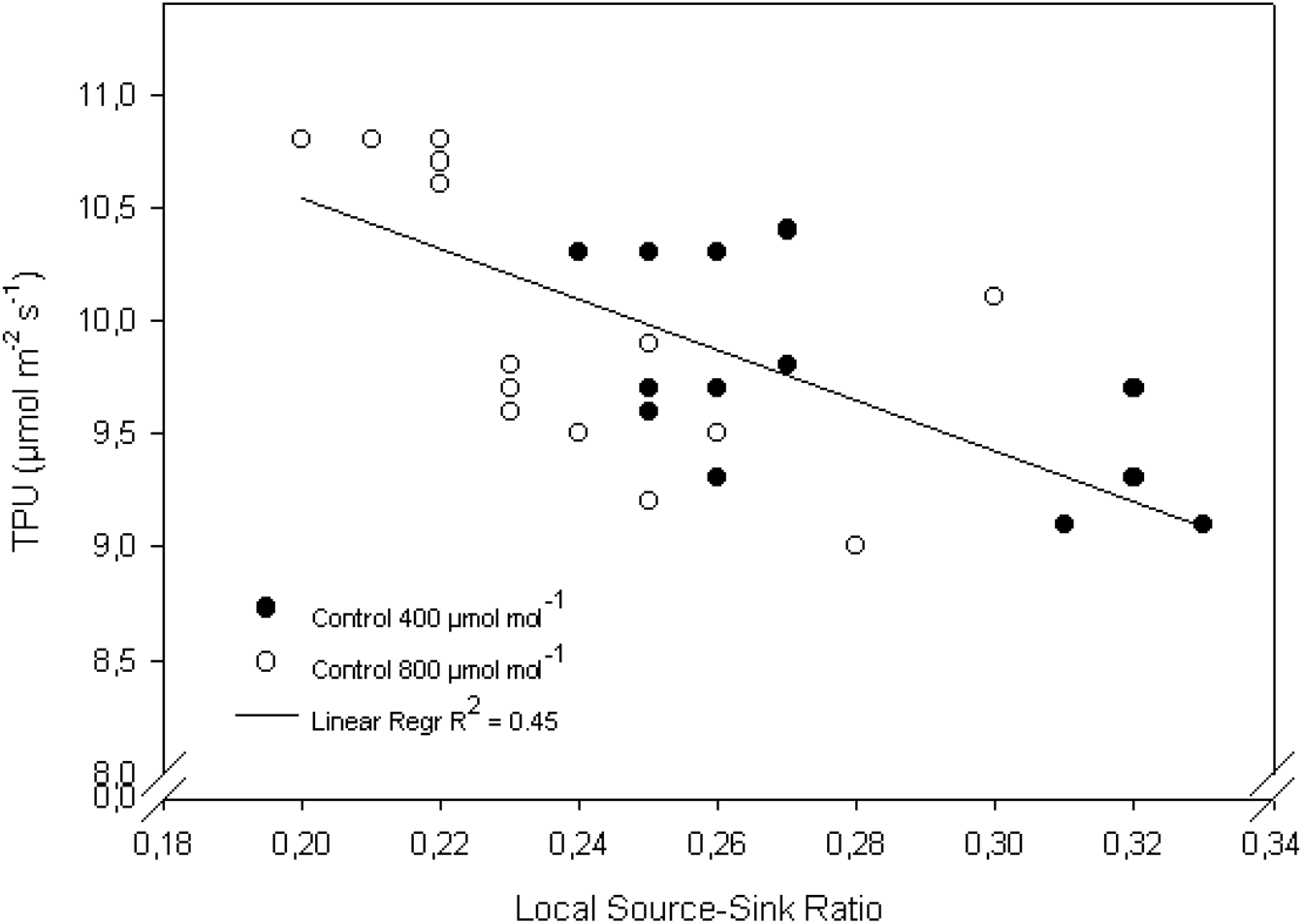
Relationship between TPU and local source-sink ratio (defined as flag leaf area / spikelet number on the main culm) in experiments 2. Line represents linear regression.

## Discussion

The physiology and biochemistry of leaf photosynthesis of major crops such as rice are well studied. So are the relations between sources, sinks and the formation of grain yield at the plant or crop scale. These processes are necessarily inter-dependent but little is known on the feedbacks causing interaction. We hypothesized that (1) source-sink imbalances are locally expressed as variations of TPU in the leaf, and (2) TPU would limit photosynthetic rate when *C*_i_ exceeds a critical level. To test the hypothesis, we manipulated the source with CO_2_ enrichment and the sink with panicle pruning. The results confirmed hypothesis (1). However, in our experiments *C*_i_ did not exceed critical levels causing TPU limitation for *A* (Hyp.2), although it came close to that level in the afternoon under combined pruning and e-CO_2_. The strong reductions of *A*, accompanied by local accumulation of assimilates, confirmed the presence of feedback inhibition of photosynthesis under sink limitation.

### Photosynthesis down-regulation under C sink limitation

Elevated [CO_2_] enhances plant C source capacity in C_3_ plants and potentially, if plant sinks are insufficiently plastic, the C source-sink ratio. Our study showed that photosynthesis decreased along the day. The extent of the decrease depended on C sink limitation induced by sink pruning and/or source stimulation with CO_2_. Declining photosynthesis along the day was previously reported under non-modified C source-sink balance (Ishihara and Saitoh, 1987; Koyama and Takemoto, 2014; Yang et al., 2008). Our observations on control plants under ambient [CO_2_] confirmed this trend, whereby enhanced source and pruned sinks further amplified it. Reductions in *A* attained 50% for both factors combined at the end of the day despite constant light resources. Sink limitation effects of this magnitude have not been noticed for rice previously. A similar effect, however, has been reported for wheat (King et al., 1967) after removing ears under ambient [CO_2_].

In the present study, CO_2_ enrichment was applied during two weeks following heading. A small but significant reduction of N content per leaf area was observed under high [CO_2_], particularly in control plants. As indicated by leaf N concentration per unit dry mass, which was 36 mg g^−1^ for the elevated CO_2_ treatment, N was above the empirical observation to affect growth, reported to be 28 mg g^−1^ (Seneweera et al., 2005: study made at 700 μmol mol^−1^ CO_2_). Leaf N concentration can decrease under CO_2_ enrichment due to dilution, causing reduced photosynthesis (Ainsworth and Long, 2005; Leakey et al., 2009; Nakano et al., 1995; Yin 2013). This was avoided in this study by limiting CO_2_ treatment to two weeks, at a stage when leaves were not expanding anymore. It was also reported that under elevated CO_2_ and sink limitation, a high leaf N content could alleviate a photosynthetic down-regulation during the day (Makino et al., 1997; Seneweera et al., 2002). In our experiments, however, a diurnal decline of photosynthesis happened in all treatments despite ample N resources.

We also investigated leaf potassium concentration in Exp2 because K deficiency potentially affects photosynthesis through stomatal responses *via* osmoregulation in guard cells (Jin et al., 2011; Wang et al., 2013; Weng et al., 2007), and also assimilate transport in phloem (Gerardeaux et al., 2010). No potassium deficiency was observed that could explain the observed variations in photosynthesis.

A reduction of stomatal conductance was observed under sink pruning treatment, as reported for many plants, e.g., citrus (Nebauer et al., 2011; Urban, 2004) and coffee (DaMatta et al., 2008). Compared to control plants, panicle-pruned plants showed a smaller increase of photosynthetic rate in response to e-CO_2_, whereby pruning always reduced stomatal conductance. However, there was no significant difference in *C*_i_ between control and pruned plants when measured at a given atmospheric [CO_2_]. Pruning thus decreased *A* at an unchanged *C*_i_ level, indicating that the photosynthetic capacity of the leaf was affected. Shimono et al. (2010) also reported that rice plants with pruned panicles under ambient and elevated CO_2_ had unaltered *C*_i_ levels.

No CO_2_ effect was observed on stomatal conductance, in contrast to reports by Ainsworth (2008) and Yoshimoto et al. (2005). In our study, a supplemental dose of N fertilizer was applied just before heading stage. This might have maintained high stomatal conductance as previously shown in rice (Shimoda, 2012; Shimoda and Maruyama, 2014). Similar to another study evaluating short-term CO_2_ enrichment effects on mesophyll conductance (Tazoe et al., 2009), no PR and CO_2_ effect was observed on mesophyll conductance g_m_, a parameter known to be sensitive to environment and estimation method (Flexas et al., 2008; Pons et al., 2009; Singsaas et al., 2004; Sun et al., 2014). In our experimental conditions involving a 14-day CO_2_ enrichment, gm was very high and thus did not limit photosynthesis. Rice generally has high g_m_ as compared with other species (van der Putten et al., 2018), probably because rice leaves have high chloroplasts coverage on the mesophyll cell periphery (Busch et al., 2013). Therefore, g_m_ was not responsible for the decline in *A* observed under sink limitation. In addition, no difference was observed for SLA, a morphological trait that can affect genotypic photosynthetic capacity (Dingkuhn et al., 1998).

The down-regulation of photosynthesis observed under elevated CO_2_ in the afternoon under C source-sink imbalance is a phenomenon that escapes observation if photosynthesis time courses are studied day-to-day and not within the day. This phenomenon is not captured in measurements at daily intervals (e.g., Makino and Mae, 1999), commonly done in the morning or at noon (Pérez-Harguindeguy et al., 2013). Our results suggest that it is crucial to capture diurnal changes of photosynthesis when studying source-sink effects on photosynthetic rate, or when estimating cumulative photosynthesis for a day.

### TPU effect on photosynthesis under sink limitation

Generally, there are transitions from one limiting factor to another, although they may be masked by a concomitant decline of several factors through feedbacks. Along *A*/*C*_i_ curves, *A* is limited by Rubisco activity (characterized by *V*_cmax_) at low *C*_i_, by RuBP regeneration (characterized by *J*_max_) at higher *C*_i_, and potentially by TPU at even higher *C*_i_. A TPU limitation is characterized by a lack of sensitivity of *A* to, or by a slight decline of *A* with, increases of CO_2_ partial pressure (Sharkey, 1985). TPU limitation can be further ascertained by a decline of φPSII with increasing *C*_i_ (Sharkey, 2016). When TPU limitation occurs, photosynthesis is affected by shortage of Pi (Paul and Foyer, 2001; Sharkey and Vanderveer, 1989) needed for ATP synthesis. In the absence of a strong C sink, TPU can be rate-limited by sucrose synthesis, thereby decreasing Pi recycling rate (Paul and Pellny, 2003). This limited regeneration is reflected in the rate at which the intermediate products of CO_2_ fixation (triose-phosphate) are converted to starch and sucrose and accumulated locally.

The patterns of diurnal decline of TPU generally mirrored those of *A* (Fig.1), resulting in a highly significant correlation between the two variables (Fig. 3). This correlation in itself, however, is not proof of a rate limitation by TPU. The *A*/*C*_i_ curves in Fig. 2A suggest that with increasing sink limitation (combinations of factors e-CO_2_, pruning, time of day), *A* tended to plateau, or even decline, at high *C*_i_ values. Chlorophyll fluorescence-based quantum yield efficiency, measured concurrently with gas exchange under exposure to 14 levels of [CO_2_], indicated the critical *C*_i_ above which TPU limited *A* for each treatment (Fig. 2B). In most of the situations studied, the critical *C*_i_ incurring TPU limitation was higher than the observated *C*_i_. In one particular situation, however (e-CO_2_, pruning, evening), the critical *C*_i_ was 363 μmol mol^−1^ and thus, similar to the observed *C*_i_ value. Consequently, although TPU decreased most strongly among the biochemical photosynthetic parameters under sink limitation, it probably did not limit *A* except, possibly, for the treatment causing the strongest source-sink imbalance.

Further studies should determine if TPU can limit *A* in a climate change context with elevated ambient CO_2_, and/or under lower temperatures to which TPU is very sensitive (Busch and Sage, 2016; Cen and Sage, 2005), and for plants having lesser phenotypic plasticity and assimilate transport capacity than rice. TPU limitations have been reported under low temperature (Sage and Kubien, 2007; Sage and Sharkey, 1987) or high CO_2_ environments (Cen and Sage, 2005; Sharkey et al., 1986). In such cases, mutual adjustment of *V*_cmax_ and TPU is observed, as these parameters decrease concurrently and in strict stoichiometry (McClain and Sharkey, 2018). Indeed, we observed a strong decrease of *V*_cmax_ at high [CO_2_] in the pruned plants (Fig. 1) concurrently with TPU, suggesting a co-adjustment. Changes in *V*_cmax_ may have contributed to the observed decrease in photosynthesis under e-CO_2_ (Makino et al., 2000; Shimono et al., 2010), possibly due to a loss of Rubisco (Long et al., 2004). It was also shown that TPU limitation activates energy-dependent quenching (*q_E_*), resulting in a deactivation of Rubisco (Sage et al., 1989; Sharkey et al., 1986). To enable photosynthesis, the carbon reduction cycle needs to regenerate RuBP, consuming ATP and NADPH produced through photosynthetic electron transport in the chloroplast. This process can be evaluated by *J*_max_ parameter (RuBP regeneration) which can also limit photosynthesis. Although we observed a significant decrease in *J*_max_ during the day in response to sink pruning, variations were too small to explain the observed variation of *A*. The two parameters correlating most with *A* were *V*_cmax_ and TPU, suggesting a tight coordination between TPU and Rubisco capacity.

Our findings suggest that TPU and *A* of rice generally decline in the afternoon, and particularly when sink is restricted. This may potentially cause overestimation of whole-day photosynthesis in crop models that do not consider rate limitations to assimilate export from the leaf (Lombardozzi et al., 2017, 2018). TPU is situated at the interface between production and consumption (or removal) of photosynthates. Thus, the mechanisms controlling this parameter can only be understood in a whole-plant context including assimilate transport and partitioning among sinkd (Yang et al., 2016).

### Sugar partitioning effect on photosynthesis and TPU regulation

Similar to our results, Morita et al., (2016), Shimono et al. (2010), Thompson et al. (2017) and Zhu et al. (2016) found that leaf sucrose concentration increased more than hexose concentrations under e-CO_2_. No increase in starch concentration in the flag leaf was observed under those conditions (Shimono et al., 2010). This can be explained by the large capacity of rice to accumulate carbohydrates in culm. Moreover, leaf starch concentration in the flag leaf (100 to 300 μg cm^−2^) was below empirical critical values (600 μg cm^−2^) reported to affect photosynthesis in rice (Weng and Chen, 1991), suggesting that leaf starch accumulation did not affect *A*.

We observed an increase in starch concentration in culm internodes in panicle-pruned plants, probably because internodes acted as alternative sinks for the panicle. Culm sucrose remobilization decreased under pruning and possibly explained the increase in starch. It may have acted as a physiological signal regulating photosynthesis as reported for sugarcane (McCormick et al., 2009; Wang et al., 2018).

Flag leaf sucrose concentration was identified as the main nonstructural carbohydrate affected by pruning treatment and [CO_2_], showing a continuous increase along the day. Photosynthesis is inhibited by leaf carbohydrate accumulation (Goldschmidt and Huber, 1992; Paul and Pellny, 2003). In this study, a negative linear relation was observed between TPU and leaf sucrose content. A theory of TPU control by Pi availability, mediated by sugar production, was proposed by Paul and Pellny (2003). According to this theory, TPU can limit photosynthetic rate through a reduced export carbon from the Calvin-Benson cycle, which in turn is related to the rate at which sugar phosphates are dephosphorylated and end-products are produced. Thus, the production and export of sucrose is essential for sustaining photosynthesis. We suggest that in our study, leaf sucrose was exported to plant sinks such as the panicle in control plants, preventing excessive build-up in the leaf. For panicle-pruned plants, sucrose could not be exported sufficiently during the afternoon. Some export occurred to the top internode, where starch concentration increased, but sucrose concentration increased in the flag leaf due to the smaller sink (Huber and Huber, 1992; Paul and Foyer, 2001). In this case, Sucrose Phosphate Synthase (SPS) feedback inhibition occurs because of an increase in the phosphorylation state of the enzyme (Huber et al., 1989). It has been shown that SPS is a substrate for SNF-1 related protein kinases, modulating SPS activity when sucrose accumulates (Sugden et al., 1999). Inhibition of sucrose synthesis may reduce export of TP from the chloroplast, causing a drop in Pi in the cytosol, leading to decrease in TPU (Paul and Pellny, 2003).

While the negative correlation between TPU and flag-leaf sucrose concentration showed a similar slope for the panicle-pruned and control plants, TPU was lower in the pruning treatment at any given sucrose concentration (reduced intercept, Fig. 4). Thus, the TPU vs. [sucrose] relationship was not the same for control and panicle-pruning treatments, and [sucrose] alone could not explain the TPU decline under C source-sink imbalance. Possibly, additional feedbacks on TPU occurred, e.g. via phloem sucrose concentration. Sucrose concentration in the leaf phloem depends on the rate of sucrose loading at the source and unloading at the sink end (Chiou and Bush, 1998; Li et al., 2003). Panicle pruning probably led to high sucrose concentrations in the leaf phloem as sucrose transport is mainly operated by phloem in rice (Regmi et al., 2016). Photo-assimilates would build up in the mesophyll (Chiou and Bush, 1998) and decrease TPU as previously described.

The ratio of flag-leaf area over the fertile spikelet number of the corresponding panicle provides a rough proxy for local C source-sink ratio during grain filling. It correlated negatively with TPU (Fig. 5), suggesting that morphology-based phenotypic plasticity causing variation in C source-sink ratio can affect TPU. A recent study also reported the effect of sink strength on sucrose partitioning that may be used to increase grain yield in rice (Morey et al., 2018). More research is needed to understand how whole-plant source-sink interactions affect crops’ ability to utilize rising CO_2_ levels.

This study provided new insights into the effect of C source-sink relationships on rice photosynthesis and in particular, its parameter TPU. A significant down-regulation of photosynthesis (up to 50%) was demonstrated during the 2^nd^ half of the day in response to sink limitation. TPU strongly decreased along with *A* and it was negatively correlated with flag-leaf sucrose concentration, suggesting sugar feedback inhibition of *A*. It is suggested that photosynthesis measurements performed in the morning, as commonly practiced, may not reliably represent the plant’s diurnal photosynthetic performance, particularly under CO_2_ enrichment or sink limitation.

Although TPU decline mirrored the decline of *A* under sink limitation, its rate-limiting effect on *A* could not be confirmed, except possibly at the end of the day for the combination of e-CO_2_ and panicle pruning. Only under these specific conditions, the observed *C*_i_ was similar to the critical *C*_i_ above which quantum yield efficiency decreased. TPU may thus play an important role in photosynthesis regulation only under extreme source-sink imbalance that may occur in plants that poorly adjust sinks and assimilate transport to increased assimilation potential.

Based on these results, it will be interesting to explore the photosynthetic responses of genotypes differing in source-sink ratio and the adaptive plasticity of sinks in CO_2_-enriched environments.

## Acknowledgments

This work was financially supported by CIRAD, French Agricultural Research Centre for International Development; http://www.cirad.fr). We thank Audrey Dardou, Paul Pruvost, Christian Chaine, Remy Michel for their valuable assistance in technical support, and the biochemical phenotyping platform of the AGAP research unit at CIRAD.

## Supplementary data

**Fig. S1:** Boxplot on various parameters for the two experiments

**Table S1:** Summary of 3-way analysis of variance of physiological flag leaf variables

**Table S2:** Summary of 3-way analysis of variance of non-structural carbohydrate concentrations

**Table S3:** Summary of 3-way analysis of variance of plant biomass and morphological variables

**Table S4:** Plant growth characteristics

**Table S5:** Type III test of fixed effects for co-variance analysis for TPU with carbohydrates and treatment

